# It takes experience to tango: Experienced cochlear implant users show cortical evoked potentials to naturalistic music

**DOI:** 10.1101/2025.06.04.657805

**Authors:** Niels T. Haumann, Bjørn Petersen, Alberte B. Seeberg, Peter Vuust, Elvira Brattico

## Abstract

Approximately 30% of cochlear implant (CI) users report that restoring their ability to enjoy music is a primary goal. However, music perception in CI users has mostly been investigated in controlled laboratory settings using simplified stimuli, such as pure tones or monophonic melodies. There is an increasing interest in developing objective measures of CI outcomes in everyday listening situations, particularly in music listening, which involves complex stimuli rich in timbre, pitch, rhythm, and overlapping sounds. One promising approach is to measure cortical auditory evoked responses (ERs) in CI users. We investigated whether ERs to sound onsets in a naturalistic four-minute music piece could be measured in adult CI users (*N*: 25; ages 18–80; CI experience: 0.3–14 years). We assumed that the accumulation of CI experience might be reflected in the morphology of the ERs. The results confirmed that P2 responses to sound onsets embedded in a whole piece of music can be detected in experienced CI users. Compared to a control group with normal hearing, the CI users showed P2 responses with lower amplitudes and longer latencies. Exploratory linear regression models suggested that the logarithmic duration of CI experience significantly predicted both perceived quality of musical sounds and P2 amplitude, explaining 38% and 28% of the variance, respectively. The findings suggest that music perception outcomes may continue to improve for up to 2–4 years post-implantation. Altogether, the results are consistent with the use of ERs to track CI adaptation to music listening.

## Introduction

For people with profound hearing loss, hearing can be partly restored with a cochlear implant (CI). Most CI users regain sufficient competency to participate in spoken conversations. However, restoring the individual enjoyment and social benefits of participating in music activities remains challenging, despite approximately 1/3 of CI users are expressing interest in regaining their ability to listen to music (Migirov et al., 2009; Kohlberg et al., 2014). The first obstacle to music perception with the CI lies in the limited ability to accurately transmit pitch, timbre, and sound intensity, due to the nature of the connection between the electrical pulses delivered by the CI and the auditory nerve fibers (for a review, see Wouters et al., 2015). There is also uncertainty about the time required for either passive adaptation or active rehabilitation to achieve optimal outcomes in music listening with a CI device (Sandmann et al., 2015; Seeberg et al., 2023). For example, a CI user describes the adaptation process for music listening in a recent qualitative study (Gfeller et al., 2019):

“*The first 3 months after activation was extremely difficult. As time went on, it became easier. About 9 months it started sounding “normal” again. I was still missing things, but my brain adjusted to the new sound, and it was becoming enjoyable again rather than effortful. Now at 1*.*5 year, it is easy and enjoyable probably 90% of the time*.”

Ongoing research and development aim at improving CI users’ perception of sounds in music, which commonly consists of pitched sounds with harmonics (i.e., integer multiples of the fundamental frequency) and non-pitched percussive sounds with broadband noise (Jiam et al., 2017; Nogueira et al., 2018). Pitched musical sounds are organized into melodies that often overlap with chords and percussive elements. To hear the individual sound onsets in the music, the listener must recognize both pitch and timbre (or spectral-temporal envelope) and detect frequency-band specific intensity increases (Alías et al., 2016; Han et al., 2016; Koning & Wouters, 2016). The perception of pitch height with CIs is primarily facilitated by electrical pulses that imitate the peaks of the acoustical oscillations, stimulating the auditory nerve fibers with shorter wavelengths peaking in the more superficial basal locations of the cochlea, and longer wavelengths peaking in the deeper apical regions (Hochmair et al., 2015). For frequencies below 300 Hz, the pitch height can also be partly transmitted by incorporating temporal fine structure from the audio waveform, whereby the electrical pulses from the CI include fine-grained oscillation cycles matching the pitch frequency (Heng et al., 2011; Müller et al., 2012; Moon & Hong, 2014). Given the complexities of the underlying biophysiological processes, it is necessary to investigate how different CI hardware and software configurations influence the final hearing outcomes.

The benefit of aided hearing in everyday listening situations is a topic of increasing interest in hearing research (Keidser et al., 2020; Beechey, 2022; Smeds et al., 2023). Field studies have been explored in everyday situations; however, because the researcher has no control over the stimuli and task variables, interpreting the results can be difficult (if not impossible) (Keidser et al., 2020). Hybrid studies offer a compromise: hearing tests are conducted in a laboratory setting where the experimenter partially controls the stimuli or tasks, as in the Common Sound Scenarios (CoSS) framework for behavioral research (Smeds et al., 2023) or the naturalistic approaches used in neuroscience research (for reviews, see Alluri & Toiviainen, 2023; Tervaniemi, 2023; Brattico & Delussi, 2024). For example, the participants can be instructed to listen attentively to music through a media device. Music perception with a CI depends on proximal cues delivered by the electrical pulse series of the CI electrodes, making it relevant to consider how well these proximal cues correspond to events in the everyday acoustical environment (Beechey, 2022). In normal hearing, acoustic events can correlate with redundant proximal cues; for instance, the event of a piano sound onset can be inferred from intensity increases across a fine-grained set of frequency bands. In degraded hearing, however, listeners rely more heavily on predictive top-down processing than bottom-up sensory input to compensate for ambiguities in the proximal cues (Beechey, 2022). According to the predictive coding theory, the central auditory system is assumed to predict unfolding musical events based on prior experience with sequences of events in music (Näätänen et al., 2017; Koelsch et al., 2018; Beechey, 2022; Vuust et al., 2022).

After the surgery in which the electrode array is inserted into the cochlea, it takes time for the brain to adjust to the new auditory cues and regain understanding of the events in the acoustic environment. Most CI users restore their ability to understand spoken language in quiet situations. A recent study found that mature post-lingually deafened CI users (age 55–85 years) achieved full outcomes in spoken language perception in quiet (measured on the NVA and LIST speech audiometry) within one year of CI experience (Claes et al., 2018). For music perception, however, an extended CI adaptation period appears necessary and seems to be accelerated through active music rehabilitation interventions (Ab Shukor et al., 2021). Nonetheless, uncertainty remains regarding the duration of the adaptation period needed to achieve optimal music listening outcomes with a CI (Sandmann et al., 2015; Seeberg et al., 2023). Adaptation outcomes can also be difficult to trace through subjective ratings and behavioral tests, particularly in children at early stages of self-report skill development (Kuki et al., 2013; Norrix, 2015; Kumari et al., 2016; Castellanos et al., 2018). One promising alternative is to trace the adaptation to the CI device by analyzing cortical auditory evoked responses (ER) measured with electroencephalography (EEG) (for a review, see Meehan et al., 2025).

Evoked responses (ERs) to sound onsets are measured as voltage peaks at fronto-central electrodes (with voltage reversals at posterior and mastoid electrode sites) (Martin & Boothroyd, 1999; Meehan et al., 2025). The P1 response is characterized by a positive peak occurring approximately 100 ms after the sound in children and 50 ms in adults (Ponton & Eggermont, 2001). The N1 is a negative peak appearing at approximately 100 ms in adults (Ponton & Eggermont, 2001; Meehan et al., 2025), typically followed by a P2 response, a positive peak around 200 ms after the sound onset (Ponton & Eggermont, 2001; Meehan et al., 2025). In CI users, larger amplitudes and shorter latencies of ERs correlate with behaviorally measured pitch change detection thresholds (*r*=.46–.74) (He et al., 2012; Liang et al., 2018; McGuire et al., 2021; van Heteren et al., 2022; Vonck et al., 2022), intensity change detection thresholds (*r*=.72–.76) (Martin & Boothroyd, 2000; He et al., 2012; Kumar et al., 2020), and speech perception outcomes (*r*=.11–.84) (Friesen et al., 2009; Liang et al., 2018; Han & Dimitrijevic, 2020; McGuire et al., 2021; Sohier et al., 2021; Vonck et al., 2021; van Heteren et al., 2022; Vonck et al., 2022; Xie et al., 2022). Abnormal morphology of the P1 or N1 response seems to indicate peripheral hearing loss in children (Ponton & Eggermont, 2001) and adults (Alain et al., 2014). Additionally, abnormally low amplitude (Ostroff et al., 2003) or delayed latency (Alain et al., 2014) of the P2 response seems to be a marker of central presbycusis in adults, a condition typically associated with degraded auditory object perception and impaired discrimination of overlapping sounds in music and speech in noise (Leung et al., 2013; Sardone et al., 2019). So far, evoked response (ER) research has focused on laboratory tests of CI users’ perception of isolated musical sounds (Sandmann et al., 2010; Torppa et al., 2012; Torppa et al., 2014; Torppa et al., 2018; Cai et al., 2020), pitch contours (Zhang et al., 2013), simplified melodies (Timm et al., 2014; Petersen et al., 2015; Petersen et al., 2020; Haumann et al., 2023a; Seeberg et al., 2023; Celma-Miralles et al., 2024), and chord sequences (Koelsch et al., 2004). However, it remains uncertain whether CI users’ ERs can be reliably measured in hybrid studies using naturalistic music stimuli from everyday listening situations.

In the present study, we adopted the naturalistic free listening paradigm to study music perception after CI adaptation. In the free listening paradigm, participants listen to real music pieces while measures of their brain activity are recorded. In the present study, we investigated whether the amplitudes and latencies of CI users’ evoked responses to sound onsets in real music might be modulated by the CI experience, the rated quality of music sounds, and the enjoyment of music. Specifically, we measured P1, N1, and P2 responses to sound onsets in a naturalistic whole music piece in a group of experienced CI users (n=15), recent (inexperienced) CI users (n=10), and an age-matched control group of normal hearing (NH) participants (n=14). All participants were adults (aged 18-85 years), age-matched between the CI and NH groups. First, we tested whether P1, N1, and P2 responses to naturalistic music could be reliably measured in each group and individual participant with a fine-tuned version of the automatic event detection function of the MIR toolbox (Lartillot & Toiviainen, 2007). We compared the ER amplitudes and latencies between CI users and NH controls, assuming that ER amplitudes would be lower and latencies longer in CI users compared to NH controls, due to the corrected but residual peripheral hearing loss and challenges in auditory object formation among CI users. Finally, we explored whether the amplitudes or latencies of the P1, N1, or P2 responses could be applied as biomarkers for tracing the development of CI experience in music listening, where the development of MMN responses to deviant tones in melodies (Seeberg et al., 2023) and spectro-temporal patterns in melodies (Celma-Miralles et al., 2024) have been reported for most of the participants enrolled in the present study.

## Materials and Methods

### Participants

Fifteen experienced CI users (median CI experience: 4.4 years, IQR: 1.0–8.3; median age: 56.9 years, IQR: 49.0–65.5) and ten recent CI users (median CI experience: 21 days, IQR: 18–32; median age: 58.5 years, IQR: 51.5–67.3) took part in the study. Prior to the test day, all CI users completed a modified Danish version of the Iowa Music Background Questionnaire (IMBQ) (Gfeller et al., 2000; Petersen et al., 2013). The experienced CI users had significantly more years of CI experience (*U*=270, *z*=4.13, *p*<.001) and rated the quality of musical sounds significantly higher (*U*=225, *z*=2.18, *p*=.029) than the recent CI users (see Table 1 for demographical and clinical details). The experienced and recent CI user groups did not differ significantly in age (*U*=186, *z*=-.47, *p*=.637), duration of hearing loss (*U*=180, *z*=-.08, *p*=.420), music knowledge (*U*=180, *z*=-.08, *p*=.420), weekly music listening hours (*U*=224, *z*=1.64, *p*=.102), or music enjoyment (*U*=203, *z*=0.39, *p*=.694). An age-matched control group of 14 participants with normal hearing (median age: 62.0 years, IQR: 59.0–66.5) also participated. The study was conducted in accordance with the declaration of Helsinki and was approved by the Research Ethics Committee of the Central Denmark Region (#55018).

**Table 1.** Demographical and clinical characteristics of the CI users. The CI users completed a modified Danish version of the Iowa Music Background Questionnaire (IMBQ) (Gfeller et al., 2000; Petersen et al., 2013). The hearing loss in years indicates the duration since degraded hearing was first detected. CI experience in days refers to the first measurement within 1½ months after switch-on. OM=Oticon Medical. AB=Advanced Bionics. The following Likert scale ranges were applied: music knowledge 1–5, music enjoyment 1–7, and quality of musical sounds 1–10. The music sound quality scores were based on averages across seven dichotomous item pairs: Doesn’t sound like music – Sound like music, Dislike – Like, Unpleasant – Pleasant, Mechanical – Natural, Fuzzy – Clear, Complex – Simple, Hard to follow – Easy to follow.

| Experienced cochlear implant users |  |  |  |  |  |  |  |  |
| --- | --- | --- | --- | --- | --- | --- | --- | --- |
| Participant ID | Age (years) | Hearing loss (years) | CI experience (days) | Device type | Music knowledge | Music listening (hrs./w.) | Music enjoyment | Music sound quality |
| CIex01 | 65-70 | 32 | 2371 | Cochlear | 2 | 2 | 5 | 6.7 |
| CIex02 | 45-50 | 36 | 3613 | Cochlear | 4 | 1 | 3 | 3.1 |
| CIex03 | 50-55 | 20 | 1621 | Cochlear | 4 | 2 | 5 | 7.1 |
| CIex04 | 55-60 | 17 | 3202 | Cochlear | 2 | 2 | 7 | 9.0 |
| CIex05 | 60-65 | 56 | 1544 | Cochlear | 5 | 3 | 4 | 8.4 |
| CIex06 | 60-65 | 10 | 5036 | Cochlear | 2 | 2 | 4 | 9.1 |
| CIex07 | 45-50 | 17 | 2793 | Cochlear | 2 | 2 | 4 | 8.1 |
| CIex08 | 65-70 | 55 | 3181 | Cochlear | 2 | 3 | 7 | 8.1 |
| CIex09 | 75-80 | 8 | 1189 | Cochlear | 4 | 2 | 2 | 3.1 |
| CIex10 | 35-40 | 9 | 381 | Cochlear | 4 | 4 | 6 | 4.9 |
| CIex11 | 35-40 | 1 | 2907 | Cochlear | 2 | 4 | 7 | 8.0 |
| CIex12 | 60-65 | 31 | 180 | OM | 4 | 1 | 1 | 1.1 |
| CIex13 | 18-25 | 17 | 370 | OM | 2 | 2 | 7 | 8.9 |
| CIex14 | 55-60 | 19 | 92 | OM | 2 | 4 | 5 | 5.0 |
| CIex15 | 65-70 | 27 | 368 | OM | 2 | 1 | 2 | 2.7 |

**Table 1. Demographical and clinical characteristics of the CI users.**
| Recent cochlear implant users |  |  |  |  |  |  |  |  |
| --- | --- | --- | --- | --- | --- | --- | --- | --- |
| Participant ID | Age (years) | Hearing loss (years) | CI experience (days) | Device | Music knowledge | Music listening (hrs./w.) | Music enjoyment | Music sound quality |
| CIre01 | 65-70 | 33 | 18 | OM | 3 | 3 | 6 | 4.9 |
| CIre02 | 70-75 | 17 | 19 | OM | 2 | 2 | 4 | 3.4 |
| CIre03 | 50-55 | 53 | 20 | Cochlear | 4 | 1 | 6 | 0.0 |
| CIre04 | 65-70 | 18 | 21 | Cochlear | 2 | 1 | 2 | 1.0 |
| CIre05 | 55-60 | 56 | 10 | OM | 2 | 1 | 5 | 3.7 |
| CIre06 | 35-40 | 18 | 20 | Cochlear | 2 | 1 | 6 | 5.6 |
| CIre07 | 30-35 | 26 | 35 | AB | 5 | 4 | 6 | 3.7 |
| CIre08 | 80-85 | 25 | 22 | Cochlear | 5 | 1 | 1 | - |
| CIre09 | 60-65 | 14 | 3 | Cochlear | 2 | 1 | 3 | 4.1 |
| CIre10 | 50-55 | 21 | 42 | Cochlear | 5 | 2 | 4 | 4.4 |

### Stimuli

All participants listened to a four-minute excerpt from a studio recording of the instrumental *tango nuevo* piece *Adios Nonino* by Astor Piazzolla (recorded in Buenos Aires 1969, from the album *Astor Piazzolla y su Quinteto*, © Circular Moves 2003). This excerpt has been validated in a previous study with a normal-hearing control group (Haumann et al., 2023b); ER results for a longer version of the same music piece have been reported elsewhere (Haumann et al., 2021). The auditory stimuli were presented in mono at a 44.1 kHz sampling rate.

### Procedure

The study was part of a larger research project, which also included the CI MuMuFe MMN paradigm (Petersen et al., 2020) and an additional study with a popular song that will be reported separately (Haumann et al., *submitted*). In the present study, evoked responses were measured while the participants listened to the 4-minute instrumental tango music piece. As previously mentioned, CI users completed a modified Danish version of IMBQ prior to the test day.

The experiment was conducted at the EEG lab facilities of Aarhus University Hospital. EEG data were recorded at a sampling rate of 1000 Hz in an electrically and acoustically shielded room using a BrainAmp amplifier system (Brain Products, Gilching, Germany). Electrodes were placed in a 32-electrode cap according to the international 10/20 system, and impedance levels were kept below 25 kΩ. Electrooculography (EOG) was recorded using electrodes placed beside and above the left eye. FCz was applied as the initial reference electrode.

During EEG recordings, participants were first presented with the CI MuMuFe MMN paradigm. After this, they were presented with the current naturalistic free listening paradigm. The order of the naturalistic stimuli was always the popular song without lyrics, the instrumental tango piece (used for the current study), and the popular song with lyrics. The total duration of the EEG recording, including both paradigms, was approximately 45 minutes.

For the naturalistic free listening paradigm, participants were instructed to listen attentively to the music piece and subsequently answer questions about their perception of the music. For all participants, the sound level was individually adjusted to a comfortable level, starting from a defined baseline of 65 dB SPL. The sound was provided to NH participants using Shure in-ear headphones in both ears. CI users received sound unilaterally, sent directly to the implant via an audio connection while keeping the microphones disabled.

### Automatic detection of sound onsets

Automatic detection of sound onsets in naturalistic music presents a methodologically complicated challenge, as the audio signal typically contains multiple overlapping voices with complex spectral-temporal patterns and background noise (e.g., Smith & Fraser, 2004; Thoshkahna & Ramakrishnan, 2008; Alías et al., 2016). An additional challenge in measuring evoked responses to sound onsets in recorded music is the brevity of the auditory cortical evoked responses, which last only a few tens of milliseconds reversing polarity, thereby requiring high temporal accuracy (Haumann et al., 2021).

To address these challenges, Brownian noise affecting low-frequency bands in the audio was suppressed using custom Matlab code. Specifically, the non-negative least squares function (*lsqnonneg*) was used to fit a 1/f curve to the average amplitude spectrum. This fitted curve served as a noise floor threshold, below which all audio was removed. To mask digital quantization errors, 90 dB white noise dither was added. Automatic onset detection was performed using finetuned functions from the Music Information Retrieval (MIR) toolbox (version 1.8.1) for Matlab (Lartillot & Toiviainen, 2007). The *mirspectrum* function was used with a 100 ms frame duration and 10 ms hop size, applying the ‘Blackmann-Harris’ window to suppress scalloping loss in the Fourier transform. The ‘Terhardt’ filter (Terhardt, 1979; Pampalk et al., 2004) was applied to simulate the effects of the human outer ear on the perceived spectra. Subsequently, spectro-temporal amplitude changes were computed using the *mirflux* function, incorporating the ‘Emerge’ filter (Lartillot et al., 2013) to minimize the influence of tremolo and vibrato. Sound onsets were then identified at peak values in the spectral flux using the *mirevents* function with the default settings. In total, 737 sound events were detected (the fine-tuned code for the automatic onset detection is available at https://github.com/nielsthaumann/nameeg).

#### EEG preprocessing

The EEG data were preprocessed using the FieldTrip toolbox for Matlab (Oostenveld et al., 2011). First, the data were downsampled to 250 Hz, high-pass filtered at 1 Hz, and low-pass filtered at 25 Hz. Unused electrodes located near the CI coil, as well as other noisy or flat-line electrodes, were replaced by interpolating signals from neighboring electrodes. The average number of interpolated electrodes was: 1.6 (range: 0–4) for experienced CI users, 1.5 (range: 0–2) for recent CI users within 1½ months, 2.3 (range: 0–4) for recent CI users at 3 months post-implantation, and 0.1 (range: 0–1) for NH controls. This interpolation was performed using FieldTrip’s *ft_channelrepair* function, which applies a default distance-weighted average from neighboring electrodes.

Eye movement and CI artifact components were isolated using the infomax independent component analysis (ICA) algorithm for EEG (Makeig et al., 1996; Delorme et al., 2007). A clear vertical eye movement component was visually identified and removed in all participants. A salient horizontal eye movement component was visually identified and removed in all CI users (except one experienced CI user) and in all NH controls. CI artifact components were also visually identified and removed based on their topographical centroids (above the implant site) and their distinguishability from auditory evoked responses and neurophysiological oscillations (Viola et al., 2011; Näätänen et al., 2017). The number of removed CI artefact components averaged 3.0 (range: 0–7) for experienced CI users, 2.2 (range: 0–5) for recent CI users within 1½ months, and 2.0 (range: 0–4) for recent CI users at 3 months post-implantation.

Following previous ER studies involving CI users, where consistent mastoid signals could not be obtained (Bishop & Hardiman, 2010; Sandmann et al., 2010; Zhang et al., 2011), EEG was re-referenced to the average across all channels. Trials were then segmented from -500 ms to +500 ms relative to detected sound onsets. Noisy trials with amplitudes exceeding ±100 *μ*V were automatically rejected. The average percentage of rejected trials was 0.3% for both experienced and recent CI users within 1½ months, 1.7% for recent CI users at 3 months post-implantation, and 0.1% for NH controls.

Subsequently, individual average EEG waveforms were computed across trials for each participant. These waveforms were then decomposed into SCA components to suppress interfering sources, including overlapping responses to closely spaced sound onsets in the music stimuli (Haumann et al., 2020).

### Statistical analysis

#### Group-level statistics

Group-level statistics were performed on fronto-central EEG electrodes (Fp1, Fp2, F3, Fz, F4, FC1, FC2, C3, Cz, C4, CP1, CP2), focusing on the time window from 30–300 ms after sound onset. To identify channels and time points that significantly deviated from the baseline (0 *μ*V) within each participant group, we applied a cluster-based permutation using a two-tailed dependent-samples *t*-test, as implemented in FieldTrip (Maris & Oostenveld, 2007). The cluster alpha threshold was set at *p*<.050 with the permutation test threshold also set at *p*<.050.

To assess differences between CI groups and the NH control group, Mann-Whitney U-tests were conducted on the individual peak amplitudes and latencies of the evoked responses (P1, N1, P2) that were statistically significant compared to the baseline (at 0 *μ*V), following individual-level statistical thresholding as described in the following section.

#### Individual-level statistics

At the individual participant level, statistically significant P1, N1, and P2 components in the SCA decompositions were identified using SCA half-split average consistency statistics (Bruzzone et al., 2021). For region-of-interest (ROI) constraints, we first limited the SCA analysis to components that peaked in amplitude at fronto-central electrodes: Fp1, Fp2, F3, Fz, F4, FC1, FC2, C3, Cz, C4, CP1, and CP2. Additionally, the P1 component was constrained to positive amplitude peaks at latencies between 30–100 ms, the N1 to negative peaks between 100–200 ms, and the P2 to positive peaks between 150–300 ms.

Individual peak amplitudes and latencies for experienced and recent CI users, as well as age-matched NH controls, were analyzed based on the average ER waveforms across the Fz and Cz electrodes, showing the highest signal-to-noise ratios in the grand average across all CI and NH groups.

#### Regression models for music sound perception with CIs

Stepwise linear regression was applied to explore potential predictors of rated quality of music sounds (scale: 1–10), as well as the amplitude and latency of ERs. The independent variables included age (years), duration of hearing loss (years), CI experience (years), music knowledge (scale: 1–5), music listening (hours per week), and music enjoyment (scale: 1–7). Significant relationships were visually inspected using scatterplots. When non-linear relationships were observed, the decimal logarithm of the independent variable was applied to linearize the relationship.

## Results

### Experienced CI users show evoked responses to sound onsets in music

The experienced CI users show significant P2 responses to the sound onsets at fronto-central electrodes (Figure 1, Figure 2, Table 2). Among the recently implanted CI users, there are no consistent evoked responses to the sound on-sets, neither within 1½ months after switch-on of the CI device nor after three months CI experience. Participants with normal hearing (NH) show P1, N1, and P2 responses to the sound onsets. The P2 responses are in the CI users compared to NH longer in latency (*U*=213, *z*=3.8, *p*<.001) and lower in amplitude (*U*=102, *z*=–2.2, *p*=.027) (Table 2).

**Table 2.**
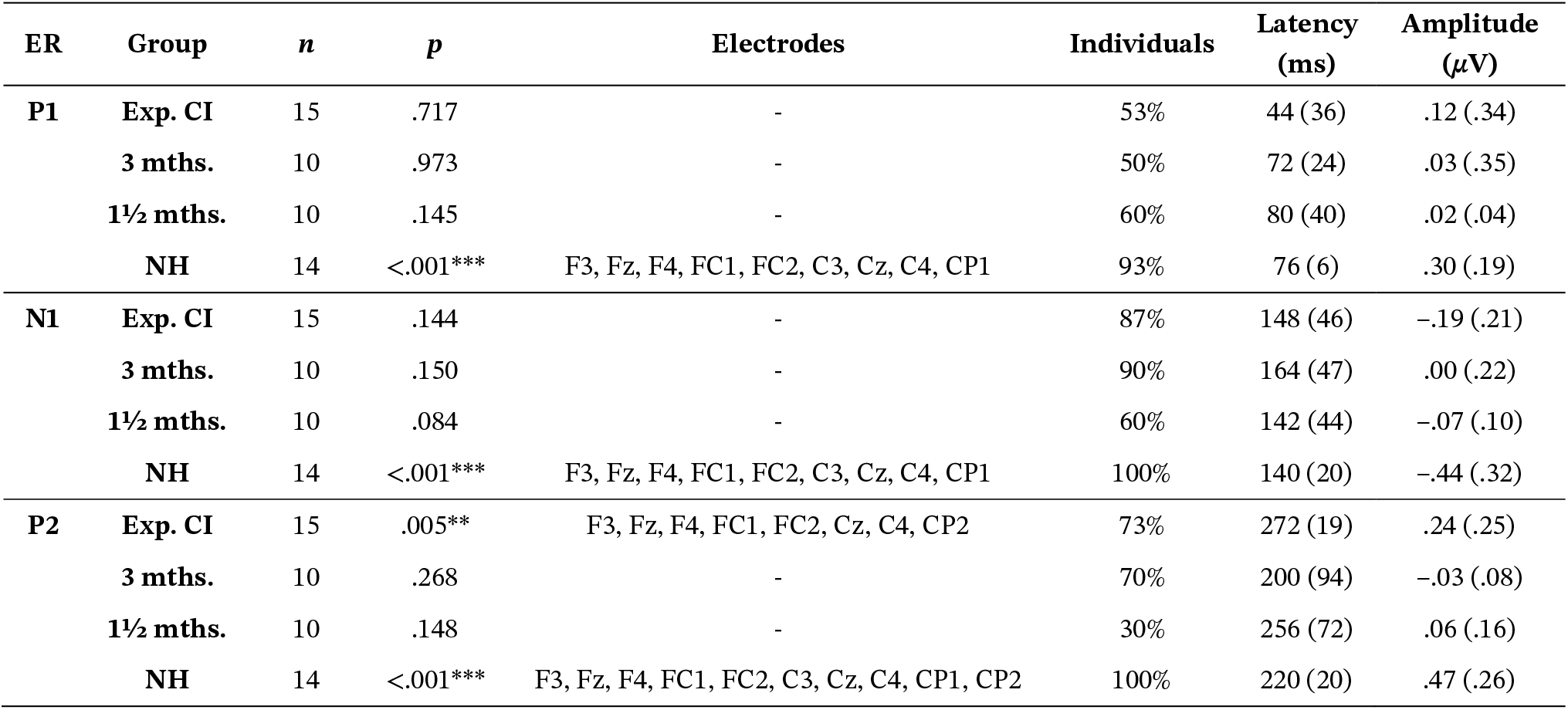
Group statistics on evoked responses. Cluster-based permutation statistics on the evoked responses (ER), P1, N1, and P2. Group: Experienced CI users, recently implanted CI users 1½ and 3 months after switch-on, and normal hearing (NH). *n*: Number of participants. The *p*-values are corrected for the number of tested clusters; ***: *p* <.001; **: *p* <.010. Electrodes: Locations are shown for clusters significantly exceeding the baseline voltage at 0 *μ*V. Individuals in percent (%) with significant ERs are shown, based on the SCA-HSAC statistic. Latency: Median latency in milliseconds (ms). Amplitude: Median amplitude in micro-Volts (*μV*). Interquartile ranges are shown in parentheses.

**Figure 1.**
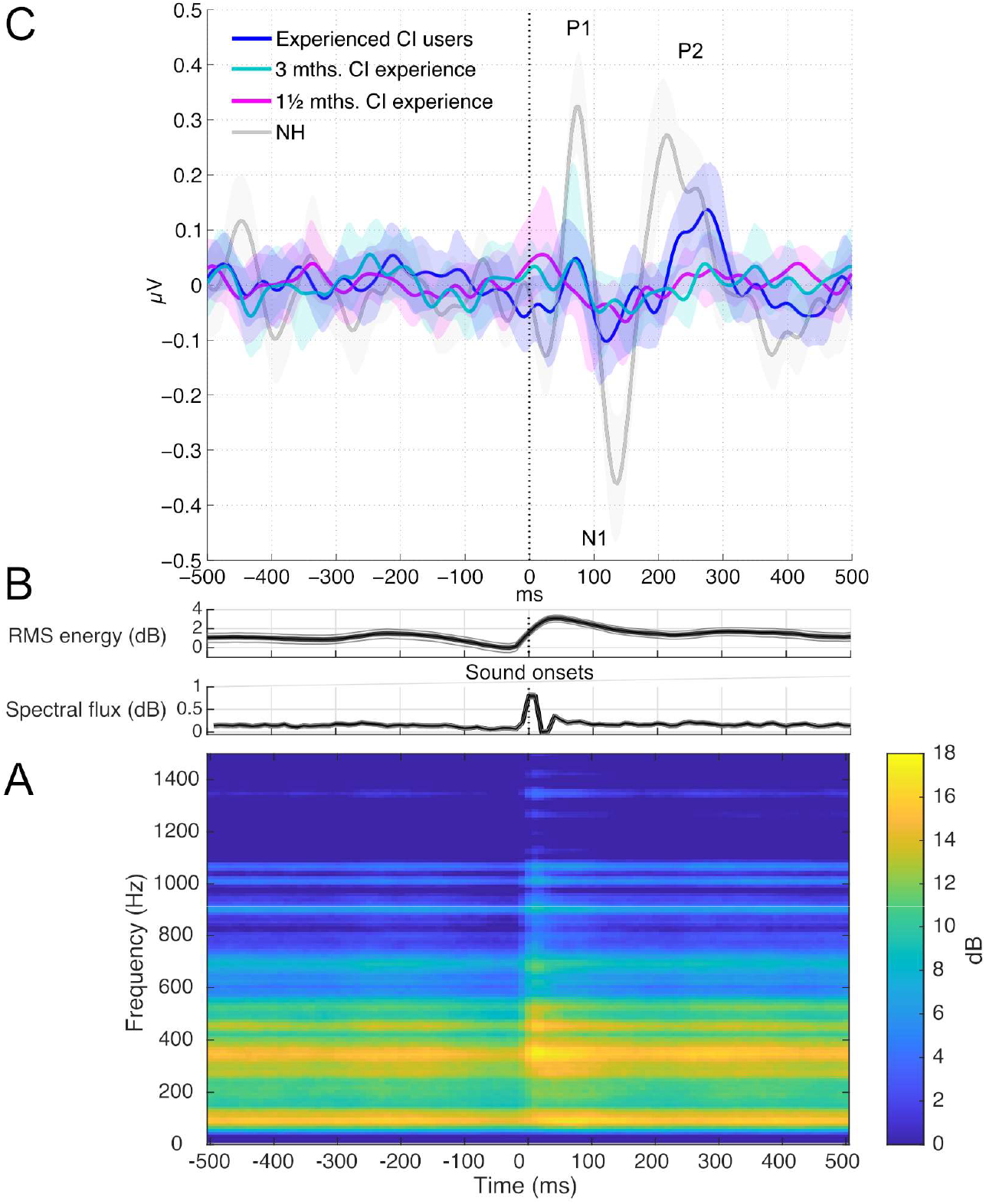
Evoked responses to sound onsets in music. **A:** The average audio spectrogram shows that the sound onsets are typically complex tones with the highest energy in frequency bands between approximately 200–600 Hz. The reference unit at 0 dB is arbitrary; the dB scale is used to show the relative change at the sound onsets. **B:** The average RMS energy and spectral flux increase at the sound onsets. C: The grand average evoked responses to the sound onsets are shown across the Fz-Cz electrodes for the experienced CI users, recently implanted CI users within 1½ and 3 months after switch-on, and normal hearing (NH). The sound onsets start at 0 ms. The shaded error bars indicate Bias-Corrected and accelerated (BCa) bootstrap 95% confidence intervals.

**Figure 2.**
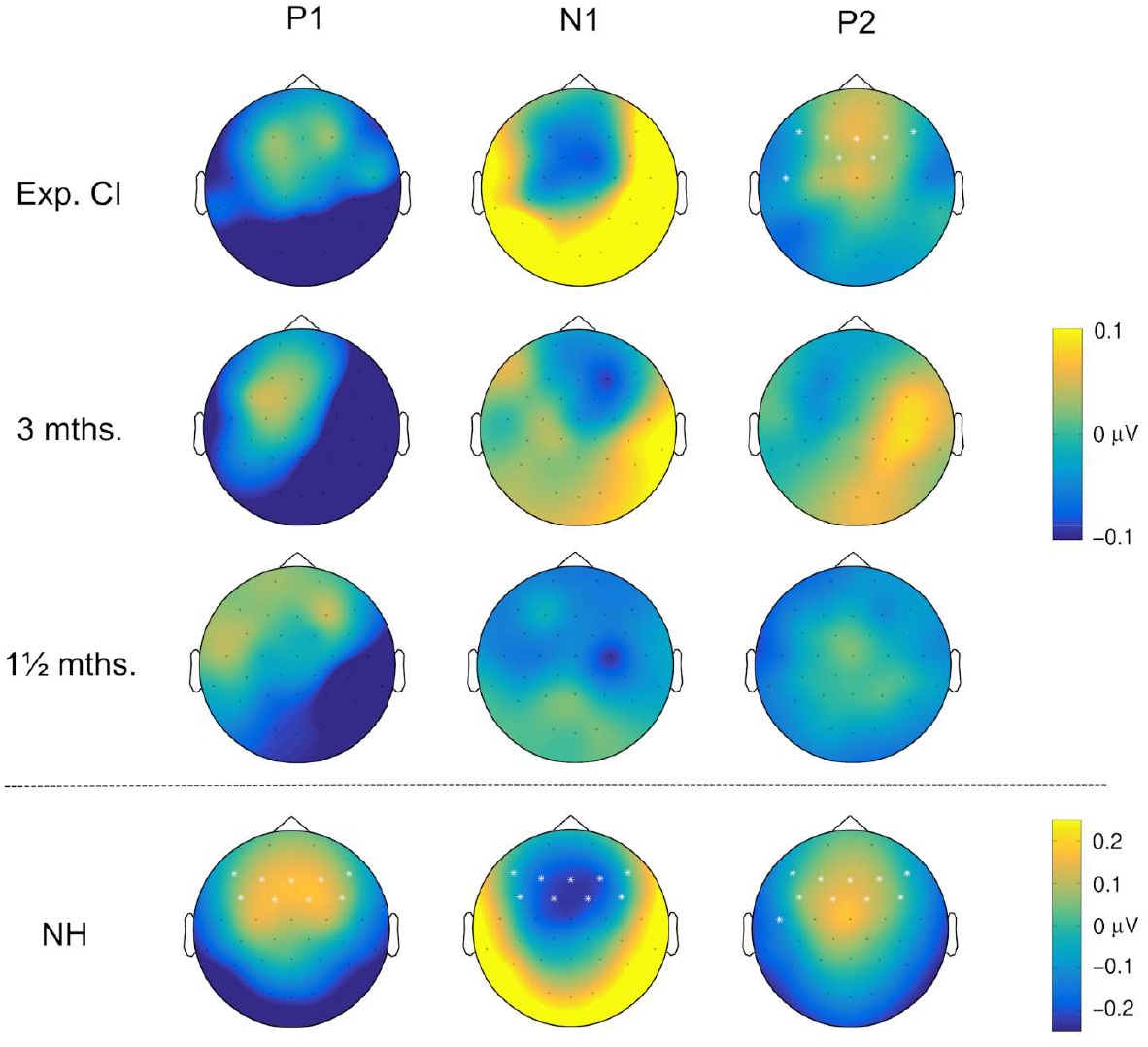
Scalp topographies of the evoked responses. The figure shows the scalp topographies across the electrodes for the P1, N1, and P2 responses based on the grand averages across the experienced CI users, recently implanted CI users within 1½ and 3 months after switch-on, and normal hearing (NH).

### Individual CI users’ evoked responses to music sounds

In the experienced CI user group, significant P1 responses are detected in 53% of the individuals, and significant N1 responses are detected in 87% of the individuals, however, the evoked responses show high variability in latency and amplitude across the experienced CI group (Table 2). Individual ERs and statistical findings are shown in Figure 3.

**Figure 3.**
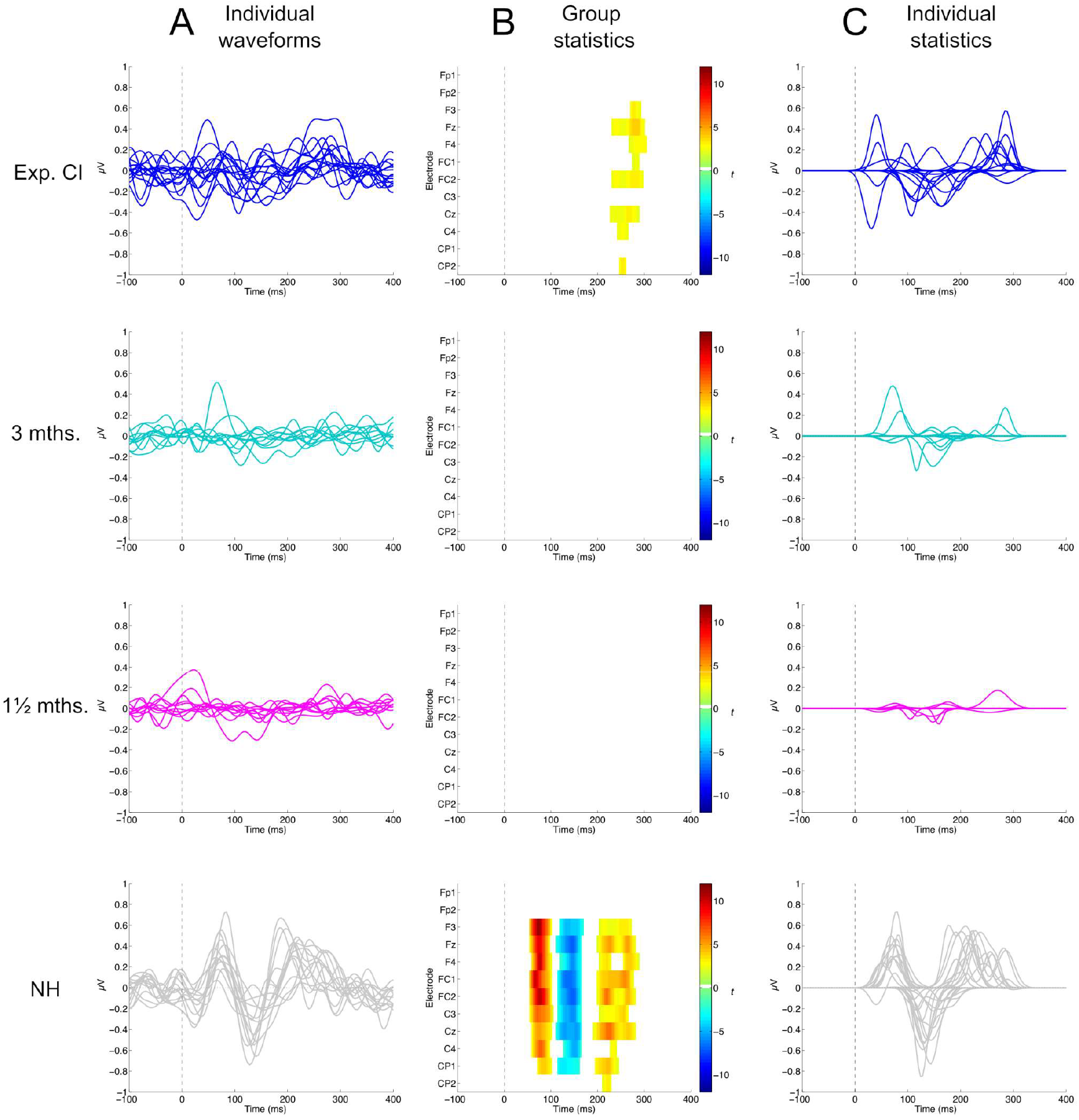
Individual statistics on the evoked responses. **A:** The individual participants’ evoked response waveforms (average across Fz and Cz electrodes). **B**: Heatmaps with *t*-values (warm colors indicate positive values, and cold colors indicate negative values) from the one-sample cluster-based permutation tests showing the electrodes and time points of the clusters with significant group responses differing from the baseline voltage at 0 *μ*V. **C**: Individual evoked response waveforms after statistical thresholding with the SCA-HSAC statistic. The statistical results are shown for the experienced CI users, the recently implanted CI users within 1½ and 3 months after switch-on, and the individuals with normal hearing (NH).

### Regression models of music sound processing with CIs

The exploratory stepwise regression models suggest that 38% of the variance in the CI users’ rated of music sounds can be explained by a logarithmic increase in days of CI experience (*R*^*2*^(25)=.38, *p*<.001), which is moderated by a linear increase in the rated enjoyment of listening to music (*R*^*2*^(25)=.25, *p*=.011) (Figure 4, top). A combined model with days of CI experience and enjoyment of listening to music explains 67% of the variance in the quality of music sounds rated by the CI users. The model predicts that the largest improvement in the music sounds occurs during the first two years of experience with the CI, which is the estimated timespan for reaching half of the maximum rating (1.78 log10(2*365) + 0.74*4 –2.54 ≈ 5, where 10 is the maximum rating, and the music enjoyment rating is adjusted to the middle value 4 of 1–7)

Similarly, 28% of the variance in the P2 amplitude can be explained by a logarithmic increase in days of CI experience (Figure 4, bottom). The rated enjoyment from listening to music does not reach statistical significance as a predictor of the P2 amplitude (*R*^*2*^(25)=.15, *p*=.054). A follow-up bivariate correlation shows that the CI users’ rated quality of music sounds does not significantly relate to the amplitude of the P2 responses (*R*^*2*^(25)=.12, *p*=.086). The model predicts that the CI users’ P2 amplitude reaches approximately half the amplitude (.24 *μ*V) compared to the group with normal hearing (.47 *μ*V) after four years CI experience (0.12 log10(4*365) –0.14 *μ*V ≈ .24 *μ*V, where .47 *μ*V is the average P2 amplitude in the individuals with normal hearing).

**Figure 4.**
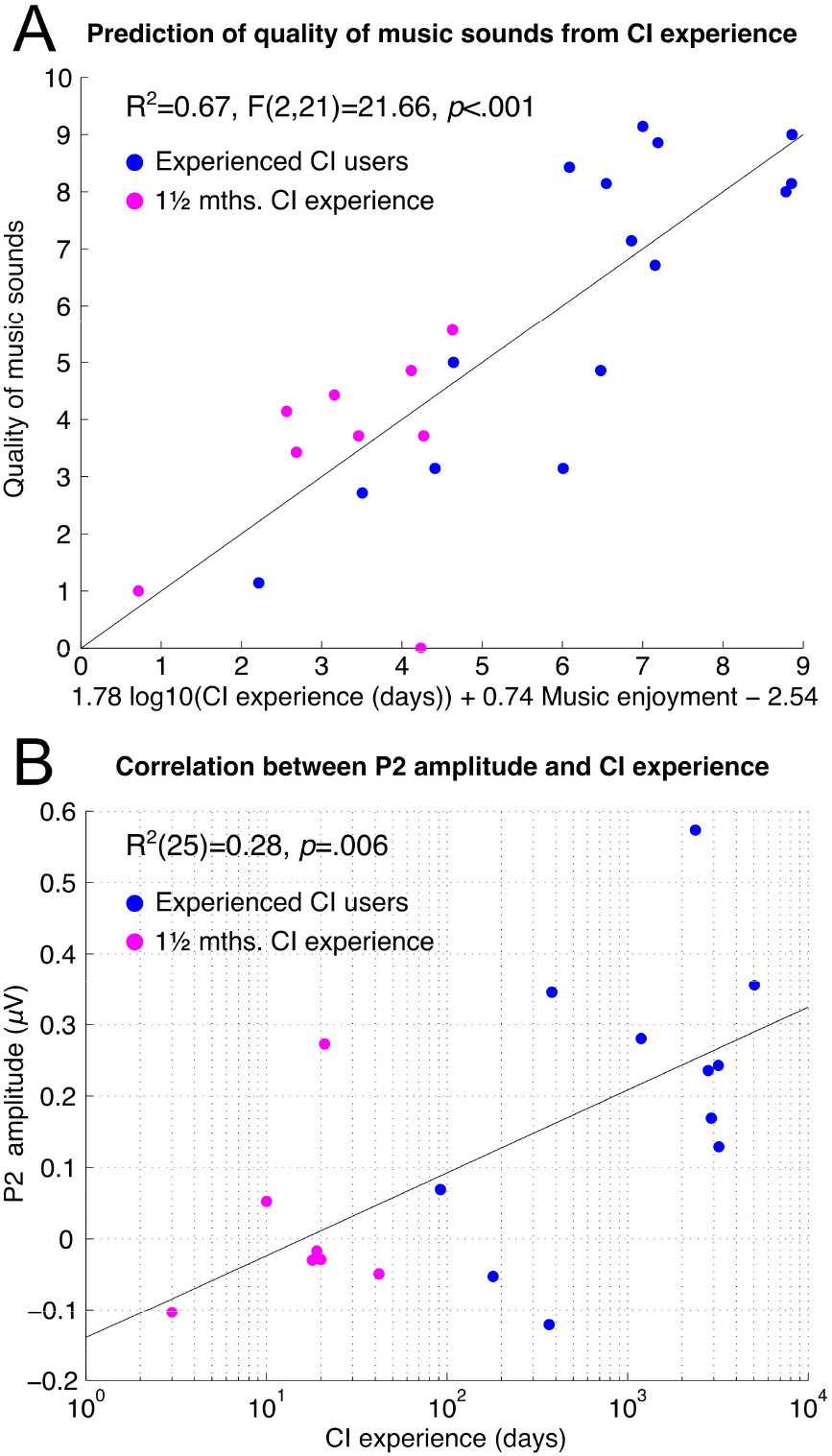
Correlational models of music sound processing with CIs. **A:** Stepwise linear regression suggests that the CI users’ rated quality of music sounds can be predicted by the CI experience (days) and music enjoyment (Likert scale between 1–7). The first two years (730 days) of CI experience seem to involve the largest logarithmic increase in the rated quality of music sounds. **B**: A positive linear relationship is observed between the P2 amplitude and the CI experience (days). With approximately four years (1460 days) of CI experience, a P2 response seems to occur in the CI users at half the amplitude (.24 μV) in comparison to the group with normal hearing (.47 μV). The magenta points indicate individuals from the recently implanted CI group measured within 1½ months after switch-on, and the blue points indicate individual experienced CI users.

## Discussion

The present study confirms that evoked responses (ERs) to naturalistic music can be measured in CI users. The P2 responses were lower in amplitude and longer in latency in CI users compared to participants with normal hearing (NH). The findings also suggest that P2 amplitude may be logarithmically related to the number of days of CI experience, with CI outcomes for music listening improving over an adaptation period until reaching a ceiling effect. Moreover, individual CI users’ ratings of the quality of music sounds showed a significant logarithmic relationship with the duration of CI experience, which, in contrast to the P2 amplitude, appeared to be partly modulated by the users’ enjoyment of music.

It is currently uncertain how long an adaptation period is needed for adjusting to the new sounds in music listening after CI activation (Sandmann et al., 2015; Seeberg et al., 2023). Previous findings using tone sweeps between 1-18 semitones (100-18000 cents) suggest that the main improvement in pitch discrimination among CI users occurs within the first three months, as reflected by increased N1 amplitudes (Sandmann et al., 2015). In the present study, CI users also listened to a four-tone piano melody with inserted deviant sounds, where improvements in pitch and timbre discrimination were observed during the first three months and were greater among experienced CI users (0.3-14 years CI experience), as indicated by higher mismatch negativity (MMN) amplitudes (Seeberg et al., 2023) and auditory steady-state responses (ASSR) (Celma-Miralles et al., 2024). However, no significant P1, N1, or P2 responses were observed at the overall group level during the first three months of CI experience. This may reflect challenges in the CI device’s ability to transmit sound intensity and spectral-temporal cues in the more complex naturalistic music stimuli compared to isolated musical sounds (Sandmann et al., 2015) and melodies (Seeberg et al., 2023).

During the fitting period between 1½ and 3 months after implantation, a few individual CI users showed increased P1 and N1 amplitude (Figure 3), potentially reflecting improved auditory nerve stimulation (Ostroff et al., 2003; Alain et al., 2014) after the adjustments of the electrical stimulation protocol (Wouters et al., 2015). Alternatively, a longer adaptation period may be required for the central auditory system to reliably detect sound onsets in naturalistic music, as suggested by increases in the P2 amplitude (Leung et al., 2013; Sardone et al., 2019) and decreases in the P2 latency (Alain et al., 2014). This interpretation is supported by the significant group-level P2 response observed only in experienced CI users. According to predictive coding theory (Näätänen et al., 2017; Koelsch et al., 2018; Beechey, 2022; Vuust et al., 2022) a likely explanation could be that the CI users needed experience with naturalistic music to form priors for the sound onsets, and the neural responses followed the development of the precision of those priors after CI experience. The regression models further suggest that music perception continues to improve beyond the initial three months of CI experience, approaching a ceiling effect, as indicated by both subjective ratings of quality of music sounds and objective P2 amplitudes.

It has recently been reported that music listening outcomes with a CI are significantly related to the quality of life after device activation (Fuller et al., 2021). CI users benefit, to some extent, from the relatively high accuracy of CIs in transmitting speech and broadband percussion sounds, as indicated by reports of greatest enjoyment when listening to music with clear lyrics and percussion (Gfeller et al., 2003). However, the recovery of pitch and timbre discrimination is critical for distinguishing social cues in speech prosody (Everhardt et al., 2020), recognizing melodies and instruments in music (Drennan & Rubinstein, 2008), and enhancing the overall enjoyment of music listening (Looi et al., 2012). As demonstrated in the current study, understanding the adaptation to pitch and timbre cues from the CI device is crucial for enhancing the listening experience to real music. In an experiment where adult CI users were asked to identify musical instruments, it was found that, unlike normal-hearing participants, CI users were unable to recognize instruments based on the attack (sound onset) and release (sound offset) of the notes (Prentiss et al., 2016). Similarly, a series of studies on melodic contour perception, distinguishing whether the pitch height increases, stays constant, or decreases (Galvin III et al., 2009), showed that only 20% of the CI users could correctly identify contours (at ≥ 80% accuracy) when synthetic tones with intervals of 3 semitones (300 cents) or smaller were used, compared to 100% of normal-hearing participants. Even for larger intervals up to 5 semitones (500 cents), fewer than 50% could reliably distinguish melodic contours. With naturalistic instrument sounds, performance worsened further, and melodic contour perception declined even more when two overlapping contours were presented – conditions common in naturalistic music. To address these challenges, ongoing research and development aim to improve pitch and timbre perception by refining techniques such as electrical current steering, enhancing fine temporal structure encoding of pitch frequencies, and by reducing ambiguity in transmitted auditory cues, e.g., through source separation and remixing of overlapping musical sounds (Nogueira et al., 2018).

In the present study, CI users’ higher ratings of the quality of musical sounds were predicted by higher ratings of enjoyment of music. However, P2 amplitude was not significantly predicted by the rated music enjoyment (*R*^*2*^(25)=.15, *p*=.054). These findings may reflect the distinction between music perception and music appreciation, or between the physical detection of sounds and the subjective pleasantness of those sounds, such as the perceived brightness or dullness in music experienced via a CI (Gfeller et al., 2008). Specifically, the results might be influenced by a negativity-bias effect in ERs, where ER amplitudes tend to be higher for unpleasant sounds compared to pleasant sounds (Norris, 2021; Tiihonen et al., 2024). In this respect, a high detection rate of sound onsets (as indicated by high P2 amplitude) could coexist with low enjoyment, if the detected sounds are perceived as unpleasant, for instance, ‘fuzzy’ or ‘mechanical’ rather than ‘clear’ or ‘natural’.

With the current investigation, we provide proof of concept for applying EEG to measure evoked responses in individual CI users, demonstrating their ability to detect sound onsets during natural music listening. We present a practical solution to the challenge of automatically detecting sound onsets in natural music with sufficient accuracy to extract evoked responses, thereby replicating previous findings obtained using manual EEG segmentation (Haumann et al., 2023b).

Future studies can build on these developments in naturalistic neuroscience by, for example, investigating post-lingual CI users’ perception of self-chosen music they enjoyed prior to hearing loss (Gfeller et al., 2005; Lassaletta et al., 2008; Friedmann et al., 2025). Of particular interest would be studying CI users’ evoked responses to specific types of sounds - such as particular music instruments, pitch ranges, or polyphonic music with overlapping voices - which could help identify which sounds in natural music are especially easy or difficult to process with a CI device (Jiam et al., 2017; Gfeller et al., 2019).

Additionally, discrimination ability profiles for individual CI users, measured via surprisal responses such as the mismatch negativity (MMN), are promising objective markers of music perception with CIs (Näätänen et al., 2017; Petersen et al., 2020; Haumann et al., 2023a; Seeberg et al., 2023). These measures should also be explored in the context of naturalistic music listening to extend CI diagnostics beyond traditional laboratory paradigms into more ecologically valid listening environments.

## Limitations

This study demonstrates that P2 responses to sound onsets in a naturalistic music piece can be measured in experienced CI users. The regression models suggest that the logarithm of CI experience (in days) explains 28% of the variance in P2 amplitude; however, this result should be interpreted with caution. We verified that the differences between experienced and recent CI user groups were not significantly confounded by age, duration of hearing loss, music knowledge, weekly music listening hours, or music enjoyment. Additionally, most CI users used the same device type, with 73% of the experienced and 60% of recent CI users using Cochlear®. Nevertheless, future studies should further explore the relationship between CI experience duration and P2 amplitude in response to sound onsets in naturalistic music, ideally through longitudinal designs tracking individuals over the first four years of CI use.

## Conclusion

The study demonstrates, with neurophysiological responses to naturalistic music listening, that adapting to music perception with a CI device requires a longer adjustment period than is typically needed for restoring spoken language ability. In line with previous research, the findings suggest that music listening outcomes continue to improve beyond the first three months following CI activation. Furthermore, the study proposes that the amplitude of the P2 response to sound onsets in real music may increase logarithmically with CI experience over time, with improvements in music listening potentially continuing two to four years post-implantation before reaching a plateau. Overall, these findings support the use of neural evoked responses measured with EEG as promising biomarkers for tracking adaptation to CI devices and monitoring the restoration of music listening in individual CI users. Especially, EEG-based measures may be valuable for assessing progress in CI users with language or attention difficulties, such as children.

## Acknowledgments

The authors would like to thank all the participants for their dedication. They also appreciate Franck Michel and Kathleen F. Faulkner for providing the programming for replacement sound processors, Monica Ipsen for her tireless efforts in recruiting participants, organizing tests, and collecting data. Special thanks to Dora Grauballe, Center for Functionally Integrative Neuroscience, for her invaluable help with bookings and managing laboratory facilities, and to Christopher Bailey for his technical assistance.

## Author contributions

Niels Trusbak Haumann conducted the analyses and wrote the first version of the manuscript. Bjørn Petersen, Peter Vuust, and Elvira Brattico contributed to the conception and design of the study and the interpretation of the results. Alberte Baggesgaard Seeberg contributed to the investigation with participant recruitment and data collection. Peter Vuust contributed with funding and manuscript revisions. All authors contributed to manuscript revisions and read and approved the manuscript.

